# Biodiversity Dimensions in Mangroves: Uncovering Interactions and Spatial Drivers in the Sundarbans

**DOI:** 10.64898/2026.03.09.710587

**Authors:** Bornali Das, Abdullah Al Asif, Shamim Ahmed, Huang Xingyun, H. A. M. Fayeem, Zawyad Bin Mostofa, Esrat Jahan Ema, Adel Mahmud Zaddary, Md Amanat Ullah, Md. Mehedi Hasan Khan, Nirmal Kumar Paul, Imran Ahmed, Swapan Kumar Sarker

## Abstract

Mangroves play a crucial role in supporting global biodiversity and ecosystem functioning, yet how their multidimensional diversity interact and respond under diverse stress conditions remains underexplored. To address this gap, using species, environmental, functional trait and forest structural data collected from the permanent sample plot (PSP) network (110 PSPs) of the world’s largest mangrove ecosystem, the Sundarbans, we answer three key questions: (Q1) How are structural, functional, taxonomic, and phylogenetic diversities interconnected? We hypothesized that these diversity components are positively correlated (H1). (Q2) What are the key environmental stressors and how the diversity components are influenced by multiple stressors? We hypothesized that these stressors negatively affect all diversity components (H2). (Q3) What spatial patterns emerge in the distributions of these diversity components? Here we hypothesized that these diversity components vary across space under changing environmental conditions (H3). Our results show that taxonomic, functional, structural, and phylogenetic diversity have varying degrees of interconnection. While taxonomic and structural diversity are strongly correlated, functional and phylogenetic diversity exhibit more independent patterns, suggesting distinct ecological processes shape each dimension. Salinity, elevation, silt, community structure and downstream-upstream gradient (i.e., upriver position) have strong influences on all the diversity components although the magnitude of the influence varies. GAM results reveal that salinity and siltation act as the primary negative drivers for most dimensions; however, functional richness and divergence show a unique positive response to salinity. Furthermore, we found that community structure and upriver position significantly influence diversity patterns, often in a non-linear fashion. Though taxonomic, structural, and phylogenetic diversity show higher values mainly in the moderate and low saline areas, functional richness shows higher values in high saline areas. Overall, our results provide strong support for all the hypotheses. Our findings highlight the importance of holistic approach integrating taxonomic, structural, functional, and phylogenetic dimensions for maintaining biodiversity and ecosystem functions in dynamic mangrove ecosystems and emphasize the need for conservation efforts that target moderate-stress zones to preserve both ecological and evolutionary diversity.

**Highlights:**

- Explored the interconnection between four dimensions of biodiversity (taxonomic, structural, functional, and phylogenetic) and how they respond to multiple stressors in the world’s largest mangrove forest.
- High salinity and siltation act as the primary environmental stressors that negatively affect overall biodiversity.
- Structural diversity is strongly related to species richness, serving as a key indicator of ecosystem health.
- Functional and phylogenetic diversity follow independent spatial patterns, promoting the need for multi-dimensional monitoring.

## 1. INTRODUCTION

Biodiversity emerges through the assembly of living organisms in biological communities and shaped by ecological and evolutionary processes (Naeem et al., 2016; Swenson, 2011). However, biodiversity is inherently multidimensional, including structural, taxonomic, functional, phylogenetic, and other interconnected aspects of life on Earth that are shaped by environmental and anthropogenic stress (Fig. 1). Taxonomic diversity (TD) is represented by species richness and various species abundance-based indices (Morelli et al., 2018) Structural diversity (SD) integrates three-dimensional forest architecture with species diversity and frequently used as a robust indicator of ecosystem health, with higher structural diversity fostering greater resilience to stress (Larue et al., 2019; Mitchell et al., 2023). SD encompasses variations in forest structural properties (e.g., diameter, height, basal area, canopy layers). Recent evidence underscores the greater importance of SD in driving above-ground biomass and productivity (Ali et al., 2016; Astigarraga et al., 2023; Zhai et al., 2024). Functional diversity (FD) represents the value, range, and distribution of species’ traits that influence ecosystem functioning (Tilman, 2001). FD has practical application in the selection of trait-diverse species for ecosystem restoration, ecosystem service enhancement, biodiversity conservation, and climate change adaptation programs (Cadotte et al., 2011; Carlucci et al., 2020). Phylogenetic diversity (PD) measures the amount of evolutionary history embodied within a biological community, providing a more comprehensive representation of biodiversity that integrates lineage divergence, evolutionary distinctiveness, and adaptive breadth (Cadotte et al., 2010). Thus, PD allows resource managers and conservation practitioners to prioritize species that represent unique lineages, and habitats that cover a wider span of evolutionary history (de Bello et al., 2021; Martins et al., 2024)

**Figure 1.**
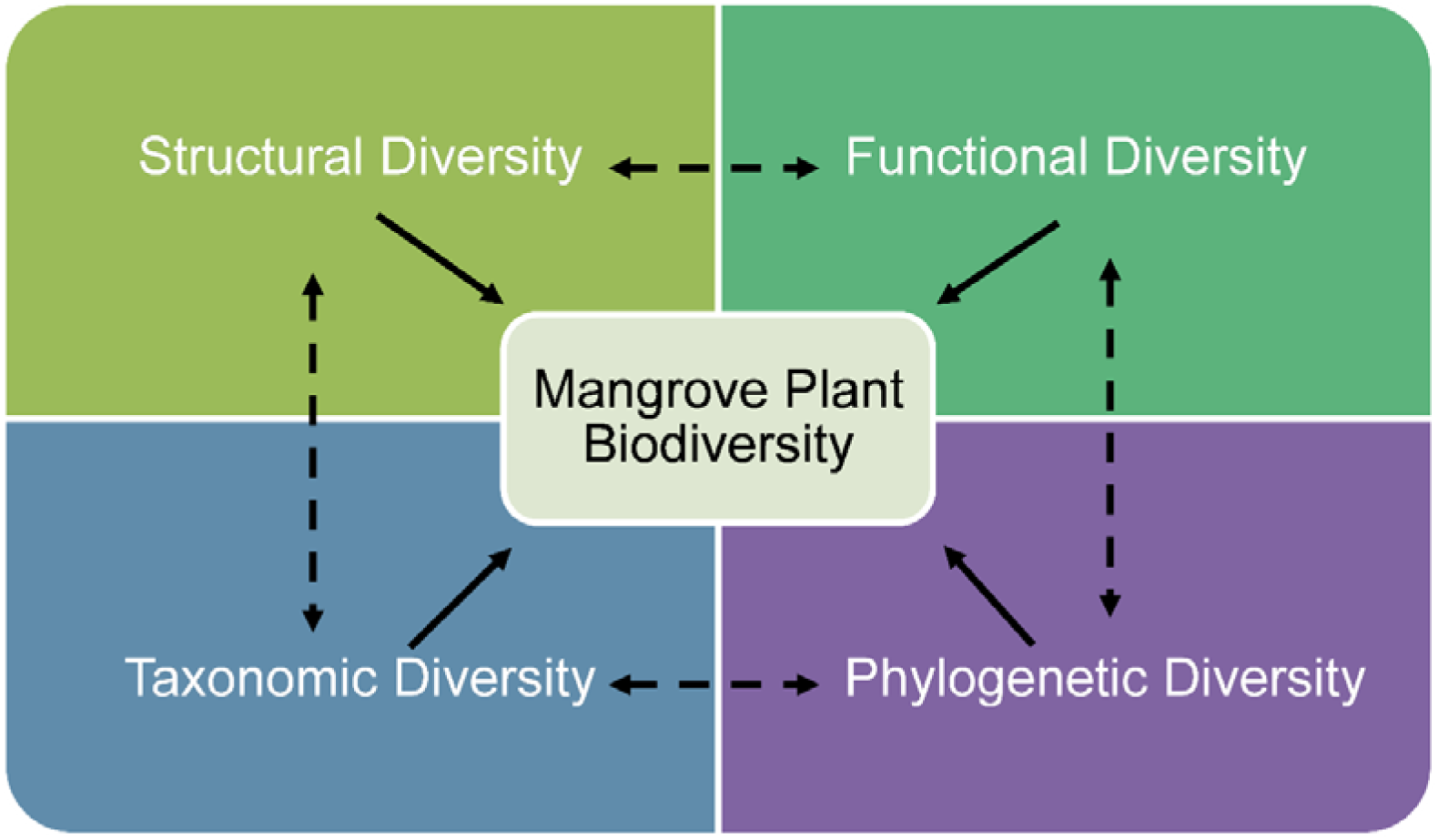
Biodiversity dimensions and their interactions.

Biodiversity indices have long been used as bioindicators or surrogates of biodiversity for measuring effectiveness of conservation initiatives (Lindenmayer et al., 2015; Rodrigues & Brooks, 2007). TD indices remain the key choice for its simplicity, ease of measurement and cost-effectiveness (Morris et al., 2014). In addition, TD indices (e.g., species richness, Shannon and Simpson diversity) are often presumed to approximate other aspects of biodiversity including FD and PD (Rodrigues, 2011). However, such over-reliance on one single measure is deemed to be misleading because not all species contribute equally to ecosystem processes and services. Further, species names and numbers ignore the function or evolutionary history of species — information vital for determining the ecological and evolutionary processes that shape biodiversity (McGill et al., 2006; Swenson, 2011). This inherent limitation of TD has raised the concern for looking at SD, FD and PD. In effect, concurrent action plans for biodiversity assessment and monitoring, forest landscape restoration, and ecosystem management strategies aim to integrate these multiple dimensions of biodiversity (LaRue et al., 2023; Wu et al., 2022) although unavailability of spatially-explicit species-environmental-trait data set has been a key challenge.

Although the multiple dimensions of biodiversity —TD, FD, PD and SD — can be interconnected, the hypothesis that one dimension is a reliable proxy for another is not theoretically or empirically guaranteed (Chapman et al., 2018; Faith, 1992; Mazel et al., 2018; Swenson, 2011). Assuming strong correlations among biodiversity dimensions can lead to incorrect inferences about unmeasured dimensions, as each contains unique ecological and evolutionary information, reflects different ecosystem aspects, and is regulated by distinct mechanisms (Pavoine et al., 2011). Although the dimensions of biodiversity are not completely independent (Naeem et al., 2025), understanding their relationships and identifying which environmental drivers regulate their spatial distributions is crucial for developing robust conservation strategies and predicting future ecosystem changes under stress (Naeem et al., 2016; Pavoine et al., 2013). However, compared to other ecosystems, such integrative approach is rarely applied to many of the vulnerable ecosystems on Earth, in particular, the coastal mangrove ecosystems (Sarker, Matthiopoulos, et al., 2019a).

Mangroves are the most productive forests in the world (Donato et al., 2011), and provide vital ecological functions and services (e.g., nutrient cycling, storm/tsunami protection, carbon sequestration, fisheries production) along the tropical and subtropical coasts (Adame et al., 2021). However, mangrove ecosystems are experiencing rapid changes in biodiversity, productivity, and ecosystem services due to sea-level rise, rising salinity, altered hydrology, coastal development, and overharvesting (Lee et al., 2021; Richards & Friess, 2016; Singh, 2020). The complex interplay of mangrove species diversity and distributions with environmental stressors (Das et al., 2026) calls for a holistic look at the multiple dimensions of biodiversity in mangrove ecosystems. However, mangrove plant biodiversity studies have rarely incorporated these dimensions, thus limiting our knowledge base on how multiple stressors concurrently impact TD, FD, PD and SD. Furthermore, there are limited understanding about the spatial variation and correlation patterns among the dimensions, which limit our ability to make evidence-based realistic conservation action plans.

Studies conducted in several non-mangrove ecosystems revealed both spatial congruence and mismatch among biodiversity dimensions, underscoring the need for multidimensional conservation strategies (Devictor et al., 2010; Socolar et al., 2016). In several mangrove ecosystem, in particular, the Bangladesh Sundarbans, environmental gradients shape clear spatial patterns in TD (Sarker, Reeve, et al., 2019), yet an comprehensive understanding of TD, FD, PD and SD and their response to multiple stressors remains limited. Clarifying the interrelationships and spatial covariation among these dimensions constitutes a critical knowledge gap.

To address this gap, we systematically examine the relationships among TD, FD, PD and SD and assess how multiple environmental drivers shape their spatial distributions in the world’s largest mangrove ecosystem, the Sundarbans, a sentinel system simultaneously impacted by climate change and human pressures. The Sundarbans Mangrove Forest (SMF), a UNESCO World Heritage Site, is a dynamic, sea-dominated delta spanning Bangladesh (6,017 km²) and India (4,000 km²). Floristically, SMF belongs to the Indo-Andaman mangrove province within the most species-rich Indo-West Pacific group and accounts for one-third of the global mangrove tree species diversity (Ghosh et al., 2015; Hossain, 2015). The detrimental effects of salinity intrusion and siltation on mangrove species distributions, species diversity, and species functional traits are well documented (Ahmed et al., 2022; Sarker et al., 2016, 2021, 2019b, 2019). Biotic homogenization has been also underway in the Sundarbans (Sarker, Matthiopoulos, et al., 2019a). The SMF is already a stressed ecosystem and projected rise in sea level (32 cm rise by 2050) (Karim & Mimura, 2008), salinity (5 – 10% decadal increase in salinity) (Mukhopadhyay et al., 2018; Wahid et al., 2007), and sedimentation (2.4 billion tons per year) (Mitra & Zaman, 2016) may further alter the biodiversity and ecosystem functioning of this vital ecosystem, thus making it an ideal system for investigating how multiple biodiversity dimensions interact with each other and how they behave under multiple stress conditions.

Using exhaustive species, traits and environmental data collected from the 110 permanent sample plots in the SMF, we aim to answer three key questions: (Q1) How are structural, functional, taxonomic, and phylogenetic diversities interconnected? We hypothesized that all these diversity components are positively correlated (H1); (Q2) What are the key environmental stressors and how the diversity components are influenced by multiple stressors? We hypothesized that all environmental stressors negatively affect all the diversity components (H2); (Q3) What spatial patterns emerge in the distributions of the diversity components? Here we hypothesized that the diversity components vary across space under changing environmental conditions and regions characterized by high taxonomic diversity also serve as hotspots for structural, functional and phylogenetic diversity (H3). Overall, this study attempts to offer a comprehensive understanding of mangrove biodiversity dimensions by tackling these issues and theories which are essential for developing pragmatic biodiversity conservation strategies (Fig. 2) and for allowing researchers to test/refine ecological theories when more idealized data sets are available in the future.

**Figure 2.**
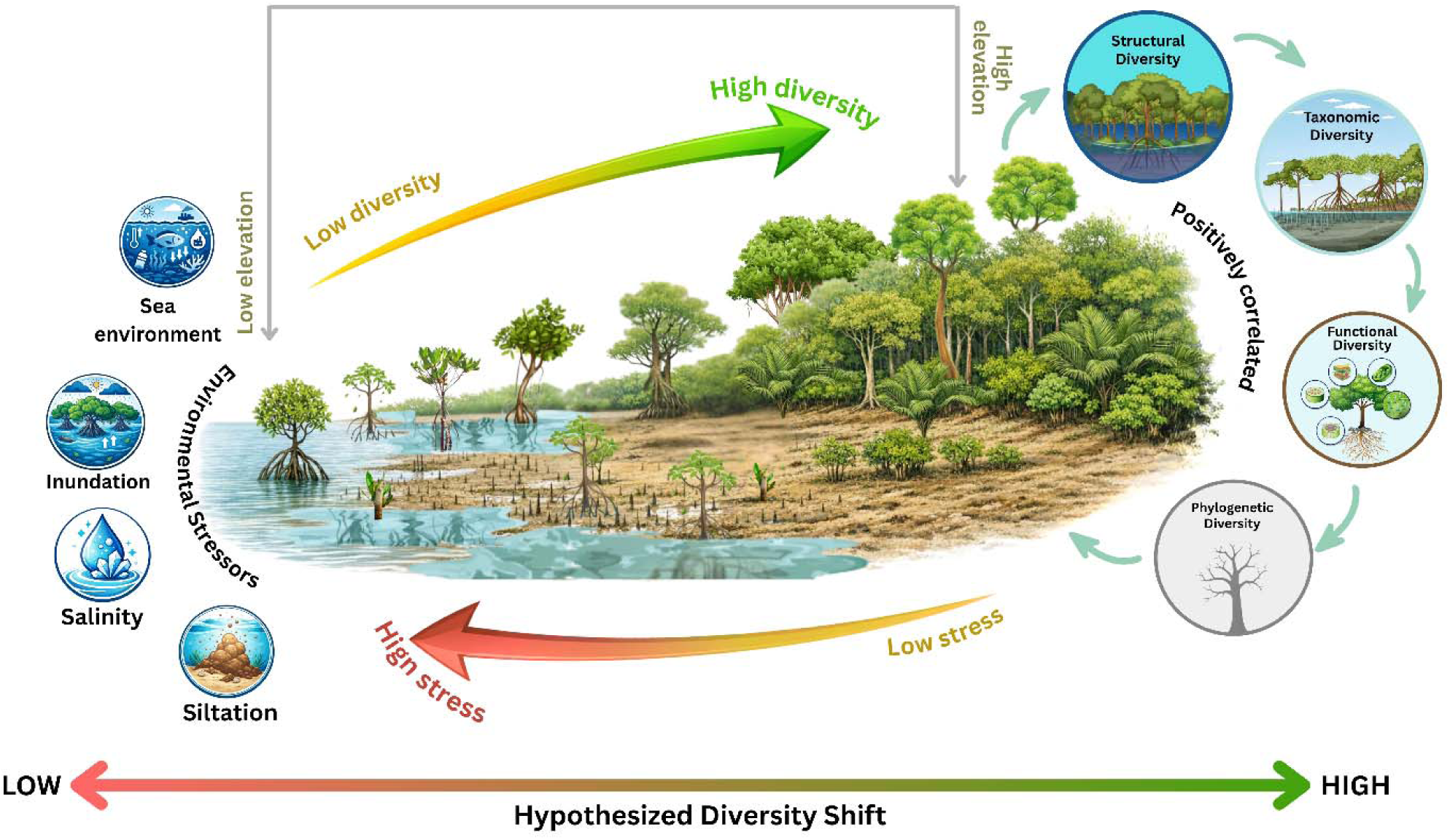
Conceptual framework for understanding the interactions between structural, functional, taxonomic and phylogenetic diversity with environmental stressors in a mangrove ecosystem.

**Figure 3.**
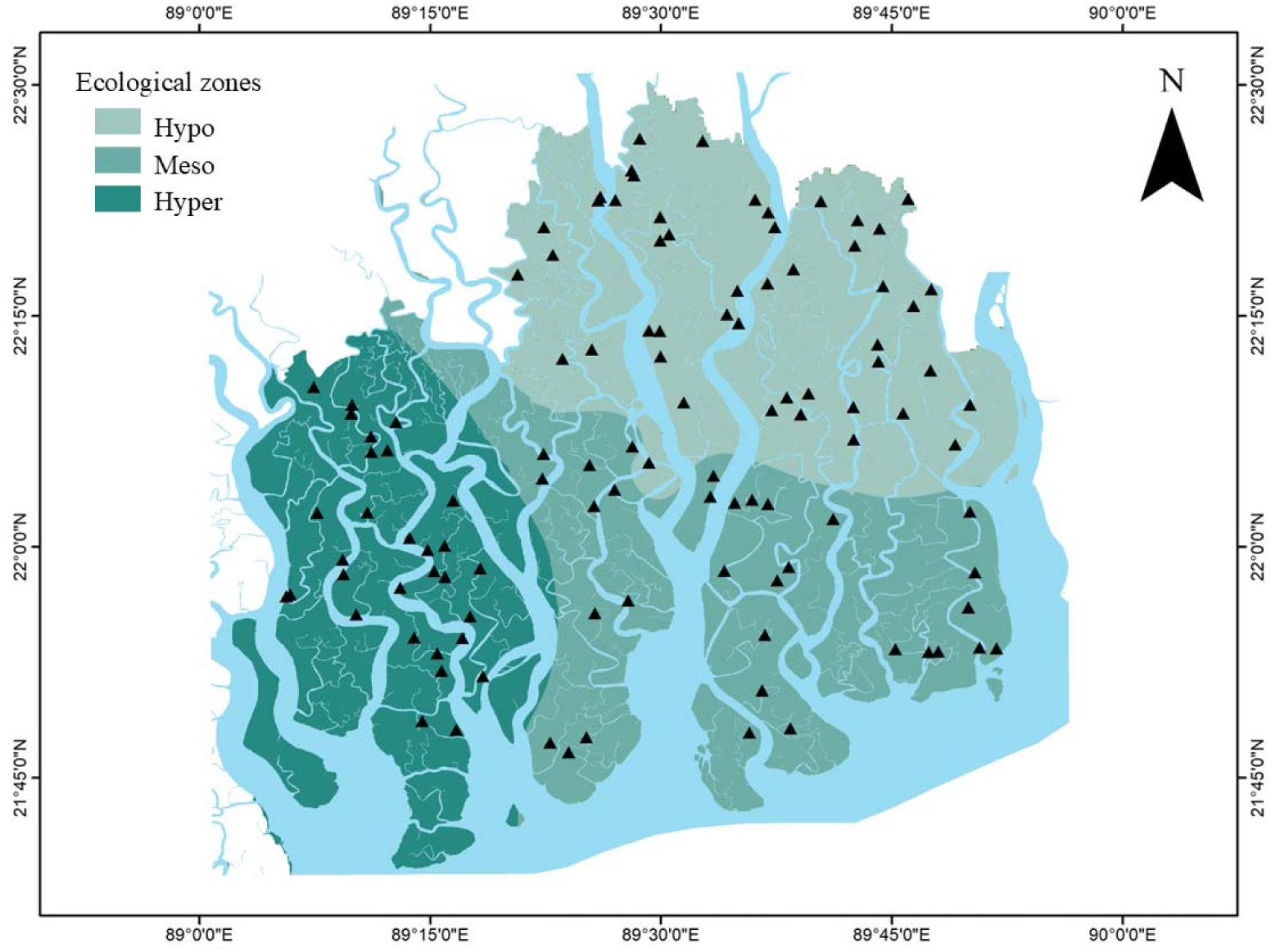
Sampling locations (black triangles) distributed in the ecological zones i.e., hyposaline, mesosaline and hypersaline zones in the Sundarbans.

## 2. MATERIALS AND METHODS

### 2.1 Study Area

The Sundarbans mangrove ecosystem covers 6017 km^2^ (21°30’’-22°30’’N and 89°.00-89°55’ E) in Bangladesh. About 69% of this forest’s topography is land and the rest are rivers, small streams, and canals. A vast portion of forest is inundated twice a day. During the monsoon, freshwater flow increases and during the dry season freshwater flow sharply drops because of the reduced water influx from the Ganges. Mandal et al. (2019) reported that the Sundarbans undergo four distinct seasons, which take place consecutively: summer, monsoon, winter, and spring. Mean annual precipitation is 1700 mm. The average temperature in pre-monsoon, monsoon, post-monsoon, and dry winter is 29°C, 30°C, 26°C and 20°C, respectively. The soil of the Sundarbans is silty clay loam with alternate layers of clay, silt, and sand. The climate is humid, maritime and tropical (Sarker et al., 2016). Based on soil salinity, Sundarbans is divided into three ecological zones, the hyposaline (<2 dS m^−1^), mesosaline (2–4 dS m^−1^) and hypersaline zone (>4 dS m^−1^) (Siddiqui, 2001).

### 2.2 Data

We collected species abundance (Table 1), structural attributes (Table 2), functional trait and environmental data from 110 permanent sample plots (PSPs) in the Sundarbans (Fig. 2). These PSPs were established by the Bangladesh Forest Department (BFD) in 1986. During the 2008 – 2014 and 2020-2022 surveys, we together with the BFD, recorded a total of 49409 trees from 20 species (Table 1). Three core functional trait data were used: Specific Leaf Area (SLA), quantified as the ratio of fresh leaf area to oven-dry leaf mass (cm² gl¹); Leaf Dry Matter Content (LDMC), calculated as the ratio of oven-dry leaf mass to fresh leaf mass (g gl¹); and Leaf Succulence (LS), defined as leaf water content per unit leaf area (g HlO cml²). Environmental variables include soil salinity (as electrical conductivity - EC), silt percentage, elevation and soil pH. To account for the influence of the downstream-upstream gradient on biodiversity components, URP (Upriver Position, the proportional distance of the PSPs from the river-sea interface) was also used as a variable. In addition, to account for the effects of species-species competition, Community Structure (total abundance of all species in a PSP) was also included in the analysis.

**Table 1.**
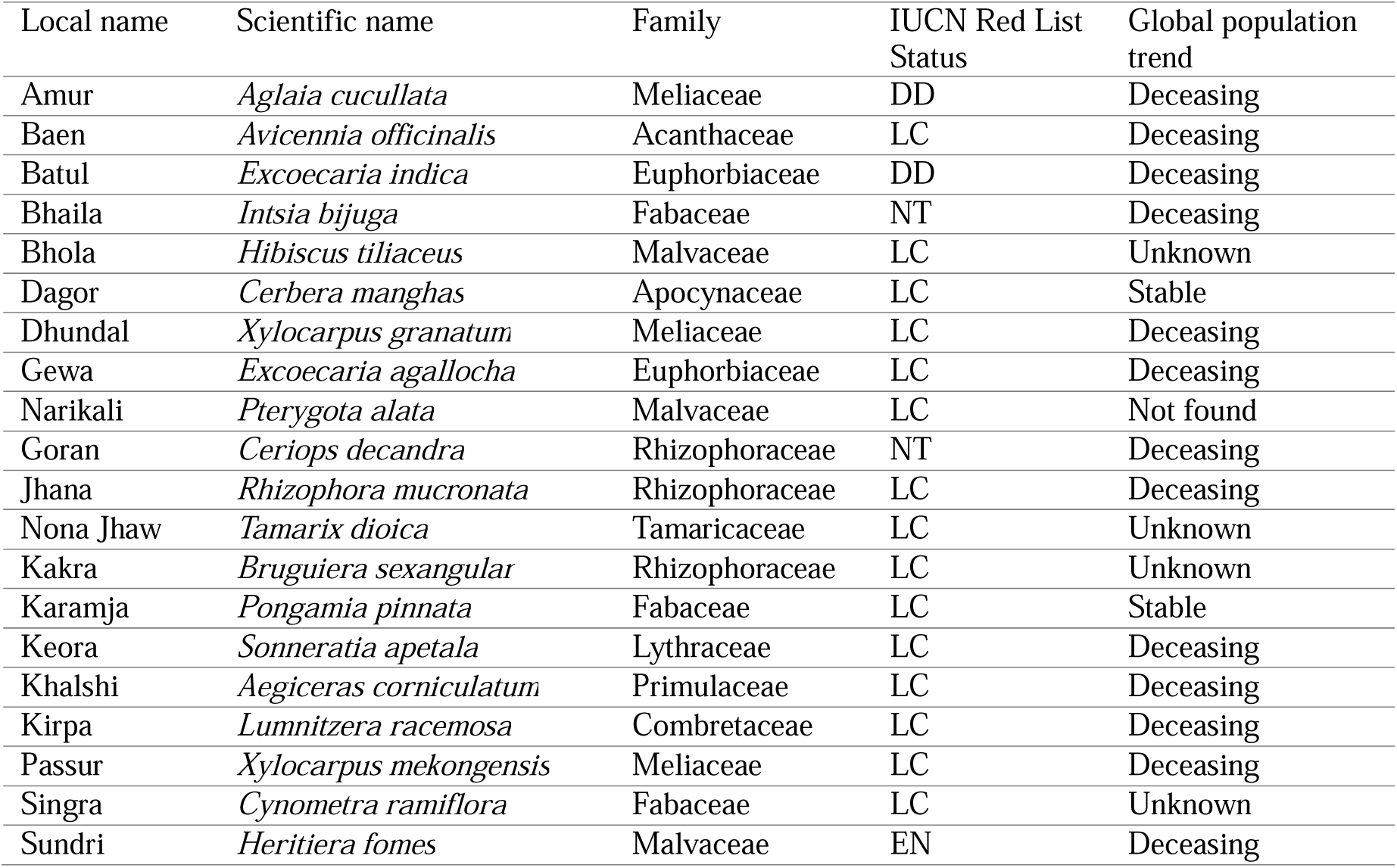
Taxonomy and global conservation status of the mangrove species censused in the 110 permanent sample plots in the Bangladesh Sundarbans. *IUCN global population trend, ^†^ Not assessed for the IUCN Red List, LC = Least concern, DD = Data deficient, NT = Near threatened, VU= Vulnerable, EN = Endangered.

**Table 2.** List of the variables used in the study.

|  | **Variables and metrics names (Abbreviation)** | **Unit/calculation** | **R Package** |
| --- | --- | --- | --- |
| **Structural Diversity Metrics** | Diameter at breast height/ (DBH) | cm | — |
|  | Horizontal diversity (coefficient of variation of DBH) (CV_DBH) | variation of DBH (cv of DBH=sd DBH/mean DBH) | dplyr |
|  | Standard deviation of tree height (SD_ht) | m | dplyr |
|  | Standard deviation of diameter at breast height (SD_DBH) | cm | dplyr |
|  | Initial total tree height (HT) | m | — |
|  | Vertical diversity (coefficient of variation of H)/ (CV_HT) | variation of H (cv of H=sd H/mean H) | dplyr |
|  | Stand density Index (SDI) | Stems ha<sup>-1</sup> | — |
|  | Canopy Packing (Canopy_packing) | Stand density index * Standard deviation of DBH | — |
| | Basal Area Per Plot (BASP) | $(\pi \times dbh^2) / 4$ | — |
| **Taxonomic Diversity Metrics** | Species Richness (Spp_Richness) | Count (number of species) | vegan |
|  | Shannon | Shannon diversity index | vegan |
|  | Simpson | Simpson diversity index | vegan |
| **Functional Diversity Metrics** | Functional Richness (FRic) | Functional richness | FD |
|  | Functional Evenness (FEve) | Functional evenness | FD |
|  | Functional Divergence (FDiv) | Functional divergence | FD |
|  | Functional Dispersion (FDis) | Functional dispersion | FD |
|  | RaoQ | Rao's quadratic entropy | FD |
| **Phylogenetic Diversity Metrics** | Phylogenetic Diversity (PD) | Faith diversity | picante |

### 2.3 Quantifying Diversity Measures

We used various tree measurements (Table 2) to assess structural diversity of the forest stands (Ahmed et al., 2025). These measurements included stand density (number of stems per hectare), mean tree height in meters, and the quadratic mean diameter at breast height (DBH), i.e., at 137 cm. For analyzing structural diversity, we calculated the coefficient of variation of H and DBH and canopy packing. To evaluate species diversity, we used three metrics: species richness (total number of different species), Shannon diversity index, and Simpson diversity index. Functional richness, evenness, divergence, dispersion, and Rao’s Q were used as measures of functional diversity (Botta-Dukát, 2005;Villéger et al., 2008). Phylogenetic diversity was quantified using Faith’s phylogenetic diversity (PD) index (Faith, 1992) which sums the minimum branch lengths on a phylogenetic tree required to include all species found for each community. To calculate the phylogenetic diversity of the species found in the study plots, a phylogenetic tree (Figure S1) was constructed and processed using R version 4.5.2. Plant species nomenclature was standardized by the plant list (http://www.theplantlist.org/). The taxonomic diversity indices were calculated using the “diversity” function of the “vegan” package version 2.7.1 (Oksanen et al. 2012). We calculated Faith’s phylogenetic diversity (PD) using the “picante” package in R (Kembel et al., 2010). Functional diversity indices were measured using “FD” package version 1.0.12.3 (Laliberté et al., 2023). FD measures several multidimensional FD indices. These indices include: functional richness, functional evenness and functional divergence. Three functional diversity indices of (Villéger et al., 2008). It also computes functional dispersion and Rao’s quadratic entropy (Botta-Dukát, 2005).

### 2.4 Statistical Analysis

In our analysis, we checked the normality of our dataset using Shapiro-Wilk normality test. To answer the first research question (Ql), we initially looked at how structural, taxonomic, functional and phylogenetic diversity correlate with each other through linear regression using the ‘lm’ function in R 4.5.2. To test whether the diversity components vary across the salinity zones we performed a Kruskal Wallis test followed by a post-hoc (Dunn) test. Several boxplots, bar plots, and other relevant applications were made from ggplot2 (4.0.2), tidyverse (2.0.0), dplyr (1.2.0), cowplot (1.2.0), multcompView (0.1-10), ggpubr (0.6.1), ggsci (3.2.0), Matrix (1.7-4), rstatix (0.7.2) etc. packages in “R version 4.5.2” (R Core Team, 2025).

We applied generalized additive models (GAMs) to identify the main stressors of the diversity components and to understand how structural, taxonomic, functional and phylogenetic diversity respond to environmental drivers such as salinity, siltation, elevation, and URP (Q2). To evaluate the influence of each explanatory variable cubic spline ‘cr’ was used and to delineate the predictor-response relationships non-parametric smoothing functions were used with effective degrees of freedom using the “mgcv” package version 1.9-3 and a global model was created (Wood, 2017). With all possible combinations of the variables, we fitted all possible candidate GAMs and the models were ranked by the Akaike Information Criterion (AIC) values using the ‘dredge’ function in the ‘MuMIn’ package version 1.48.11 (Burnham & Anderson, 2002). The relative contribution of each variable in the best-fitting model was evaluated by computing AIC values and the difference in total deviance explained between the entire model and the model with variables removed. To identify the key variables, we used the ‘importance’ function in the “MuMIn” package version 1.48.11 by determining strength of the covariates and ranked them based on their Relative Importance (RI). RI values range from 0 to 1, where 0 indicates that the target covariate is excluded from all competing models while 1 indicates inclusion in all competing models. To assess models, we also measured the goodness-of-fit of the models using the R^2^ (coefficient of determination) statistic between the observed and estimated abundance values. All statistical analyses were done in R version 4.5.2.

### 2.5 Spatial analysis

To answer Q3, we conducted spatial analysis using ArcMap 10.5. We interpolated the structural, functional, taxonomic, and phylogenetic metrics using the Ordinary Kriging approach (Li & Heap, 2014; Lian et al., 2022). Subsequently, we analyzed the interrelationships among the metrics of the four diversity types through linear regression using the ‘lm’ function.

## 3. RESULTS

At first, we examined how mangrove structural, functional, taxonomic, and phylogenetic diversity metrics varied across the three ecological salinity zones of the Sundarbans (Fig. 4) to provide an ecological context for the diversity patterns along the key environmental (salinity) gradient. These patterns reflect how different levels of salinity influence mangrove communities. For instance, reduced species richness and taxonomic diversity in hypersaline zones may result from physiological stress that filters out less tolerant species. In contrast, higher functional divergence and RaoQ in those zones suggest that a few stress-tolerant species maintain broad functional differences to survive. This descriptive section (Table S1) helps to contextualize the upcoming hypothesis-driven analyses by showing baseline patterns in the data.

**Figure 4.**
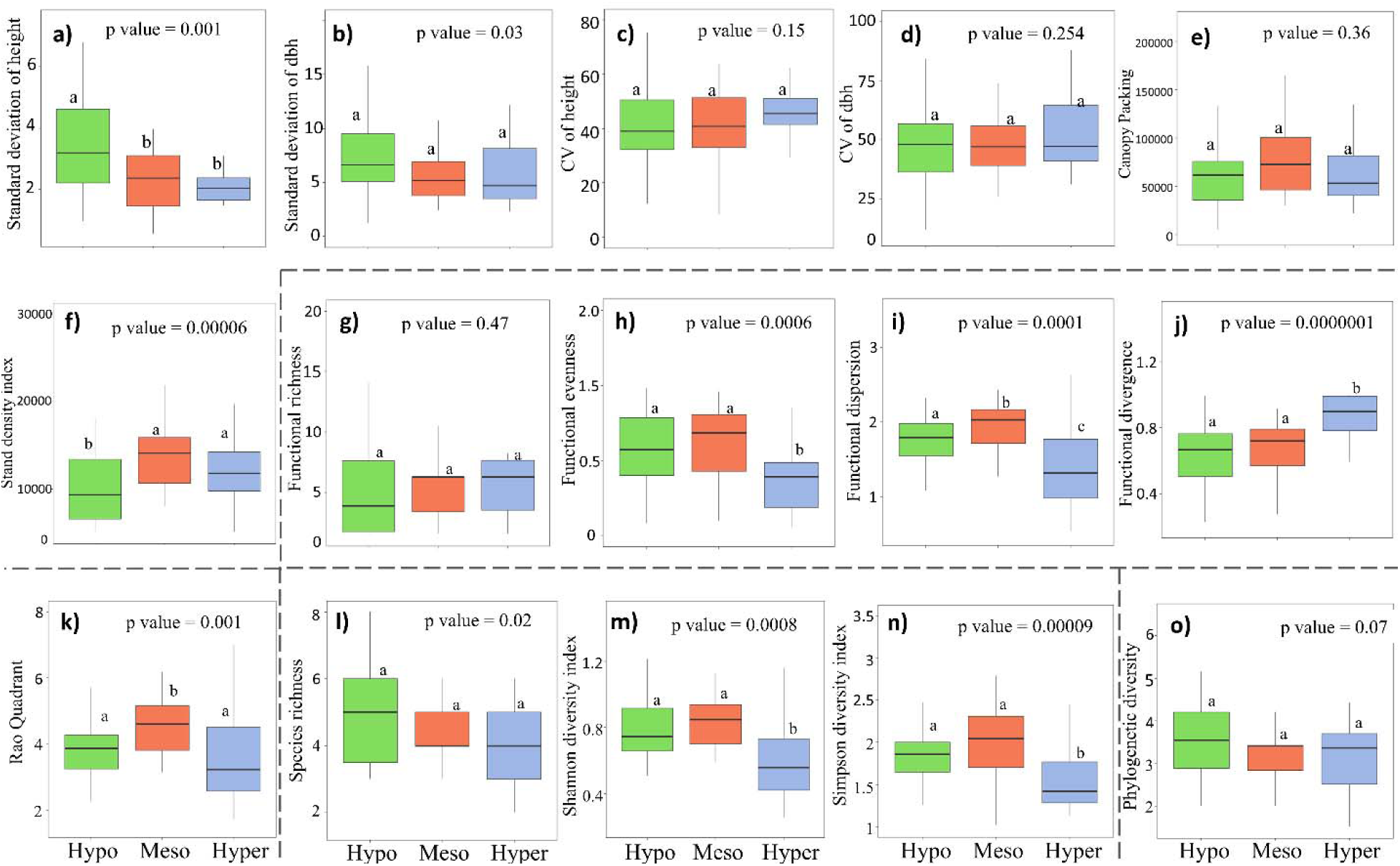
Variation in structural (a–f), functional (g–k), taxonomic (l–n), and phylogenetic (o) diversity metrics as ecological indicators across three salinity zones (hypo, meso, and hypersaline) of the Sundarbans. Boxplots show the distribution of each metric within zones. Different letters indicate significant differences among salinity zones based on post-hoc comparisons (p < 0.05).

### 3.1 Variation in Mangrove Structural, Functional, Taxonomic and Phylogenetic diversity across the salinity zones (Descriptive Results)

Significant variation in structural, functional, taxonomic, and phylogenetic diversity was observed across the salinity zones of the Sundarbans. Structural diversity metrics such as Standard deviation of tree height (*p* = 0.0016), Standard deviation of DBH (*p* = 0.03), and stand density index (*p* = 0.00006) varied significantly (Fig. 4). Overall, the hyposaline zone showed higher variability in height and DBH but lower values in stand density and canopy packing, whereas the mesosaline zone revealed the highest stand density with the hyposaline zone showing the highest height variation and the meso- and hypersaline zones exhibiting greater density. In terms of functional diversity, functional dispersion, divergence, and RaoQ increased significantly with salinity (*p* < 0.001). However, functional richness and functional evenness showed no significant differences. Taxonomic diversity declined along the salinity gradient. Species richness was highest in the hyposaline zone (*p* = 0.02), while Shannon and Simpson indices decreased significantly in hypersaline zones (*p* < 0.001). Although not statistically significant (*p* = 0.07), phylogenetic diversity (PD) showed a slight increase in the hypersaline zone.

### 3.2 Relationships between diversity components (Q1)

Overall, structural diversity (SD) metrices showed positive relationships with taxonomic diversity (TD) indices although the strength of the relationships was substantially higher with species richness (accounts for species presence-absence only) compared to Shannon and Simpson diversity indices (accounts for species relative abundance) (Fig. 5). SD metrices also showed a moderate positive relationship, with functional diversity (FRic, RaoQ) and phylogenetic diversity (PD). On the other hand, stand density index (SDI) showed a moderate negative relationship with taxonomic diversity, functional diversity (except FRic, FDiv) and phylogenetic diversity indices.

**Figure 5.**
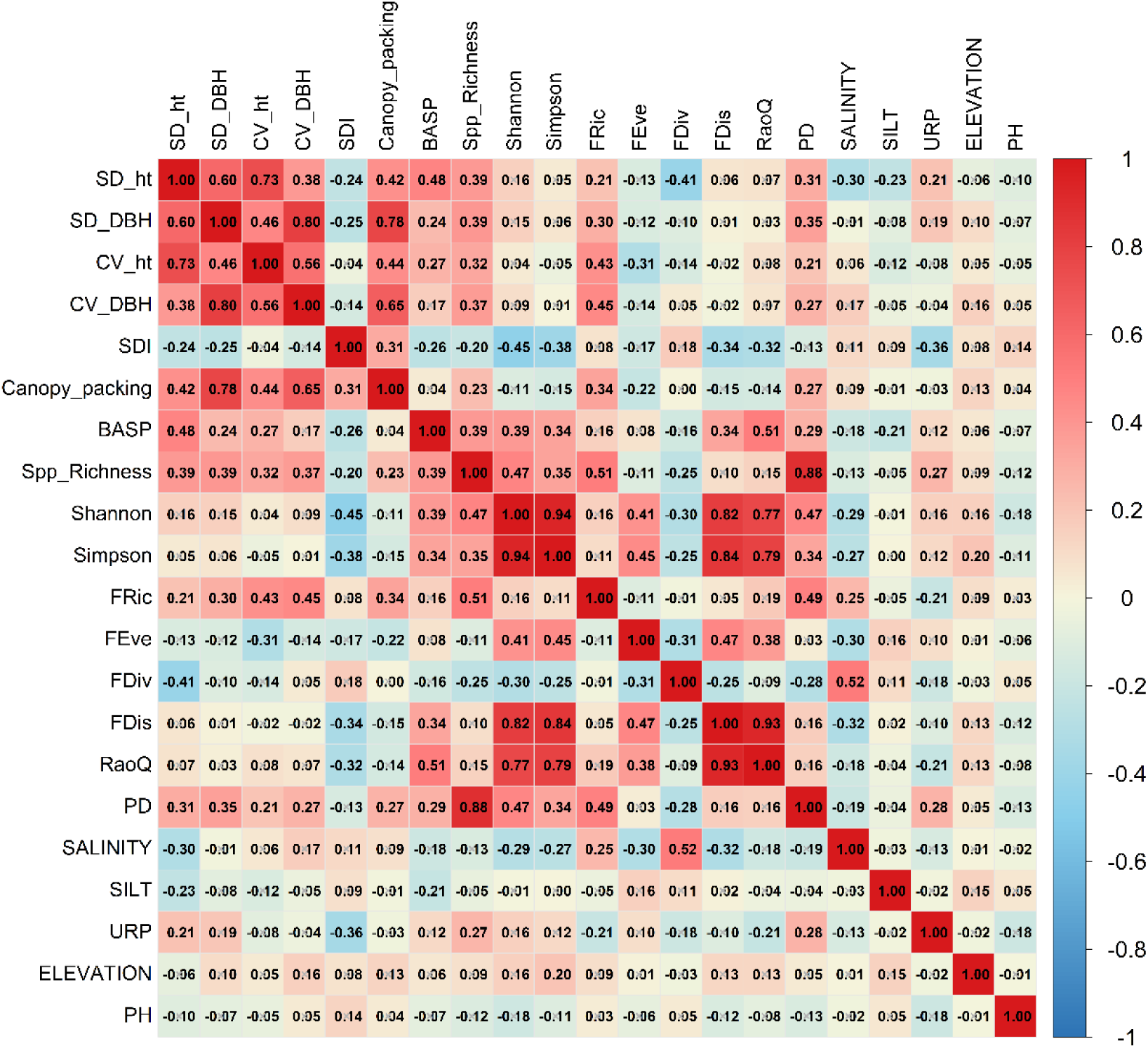
Pearson correlation matrix between streesors and structural, functional, taxonomic, and phylogenetic diversity variables. Correlation strength is represented by both color intensity and circle size, with red indicating positive and blue indicating negative correlations. This figure supports Hypothesis I and II (HI,HLJI). Abbreviations: SD_ht = standard deviation of height; SD_DBH = standard deviation of diameter at breast height; CV_ht = coefficient of variation of height; CV_DBH = coefficient of variation of diameter at breast height; SDI = stand density index; BASP = basal area per plot; Spp_Richness = species richness; FRic = functional richness; FEve = functional evenness; FDiv = functional divergence; FDis = functional dispersion; PD = phylogenetic diversity.

All taxonomic diversity indices showed a significant positive relationship with functional diversity and phylogenetic diversity indices with highest positive correlation between species richness and FRic (*r*=0.51) and, species richness and PD (*r*=0.88) and the strength of the relationships sharply dropped when we looked for the relationships between functional and phylogenetic diversity indices with Shannon and Simpson diversity. In addition, functional diversity metrices showed a moderate positive correlation with phylogenetic diversity with highest positive relationship between FRic and PD (*r*=0.49).

Among environmental variables, salinity showed consistent negative correlations with multiple structural, taxonomic, and functional diversity metrics, including SD_ht, Shannon, Simpson, FEve, and FDis, while FRic and FDiv showed positive associations with salinity. Silt content was negatively correlated with SD_ht and basal area per plot (BASP). Upriver proximity (URP) showed negative correlations with SDI and RaoQ, while PD showed a weak positive association with URP. Elevation and soil pH exhibited weak or non-significant correlations with most diversity metrics, with elevation showing a slight positive association with Simpson diversity. Overall, the correlation patterns indicate that salinity exerts the strongest influence on biodiversity metrics, with other environmental factors showing comparatively weaker associations.

While the correlation matrix provided an initial understanding of linear associations between environmental stressors and diversity metrics, it was limited in capturing potential nonlinear or threshold responses. Many ecological relationships, particularly under stress gradients such as salinity or siltation, are known to be nonlinear in nature. Therefore, to better understand how diversity patterns change across the full range of stressor gradients, we applied Generalized Additive Models (GAMs). This approach allowed us to explore more complex and ecologically realistic relationships, beyond what pairwise correlations could reveal.

### 3.3 Influence of stressors on structural, functional, taxonomic and phylogenetic diversity (Q2)

The explanatory power and the goodness-of-fit of the taxonomic, functional, phylogenetic and structural diversity GAMs varied (Table 3). Among the taxonomic diversity (TD) GAMs, Shannon diversity explained more deviance (DE = 36%) and showed better fit (Adj. *R*^2^ = 0.32) compared to those for species richness and Simpson’s diversity, suggesting the model with a moderate focus on species relative abundance in the mangrove communities could capture more signal than the models that only account for species presence-absence (species richness) or provide more importance to the more dominant species (Simpson’s diversity). Among the functional diversity (FD) GAMs, Rao’s Quadratic Entropy (RaoQ) explained more deviance (DE = 34%) and showed better fit (Adj. *R*^2^ = 0.30) than the other FD GAMs. The best phylogenetic diversity (PD) GAM explained 16% of the deviance with a relatively low fit (Adj. *R*^2^ = 0.10). Among the structural diversity GAMs, BASP, SD of height and canopy packing performed better while CV of height and SD of DBH were the least performing models, indicating within community variability in tree heights and between-plot variability in basal area and canopy packing can be better predicted in mangrove ecosystems.

**Table 3.**
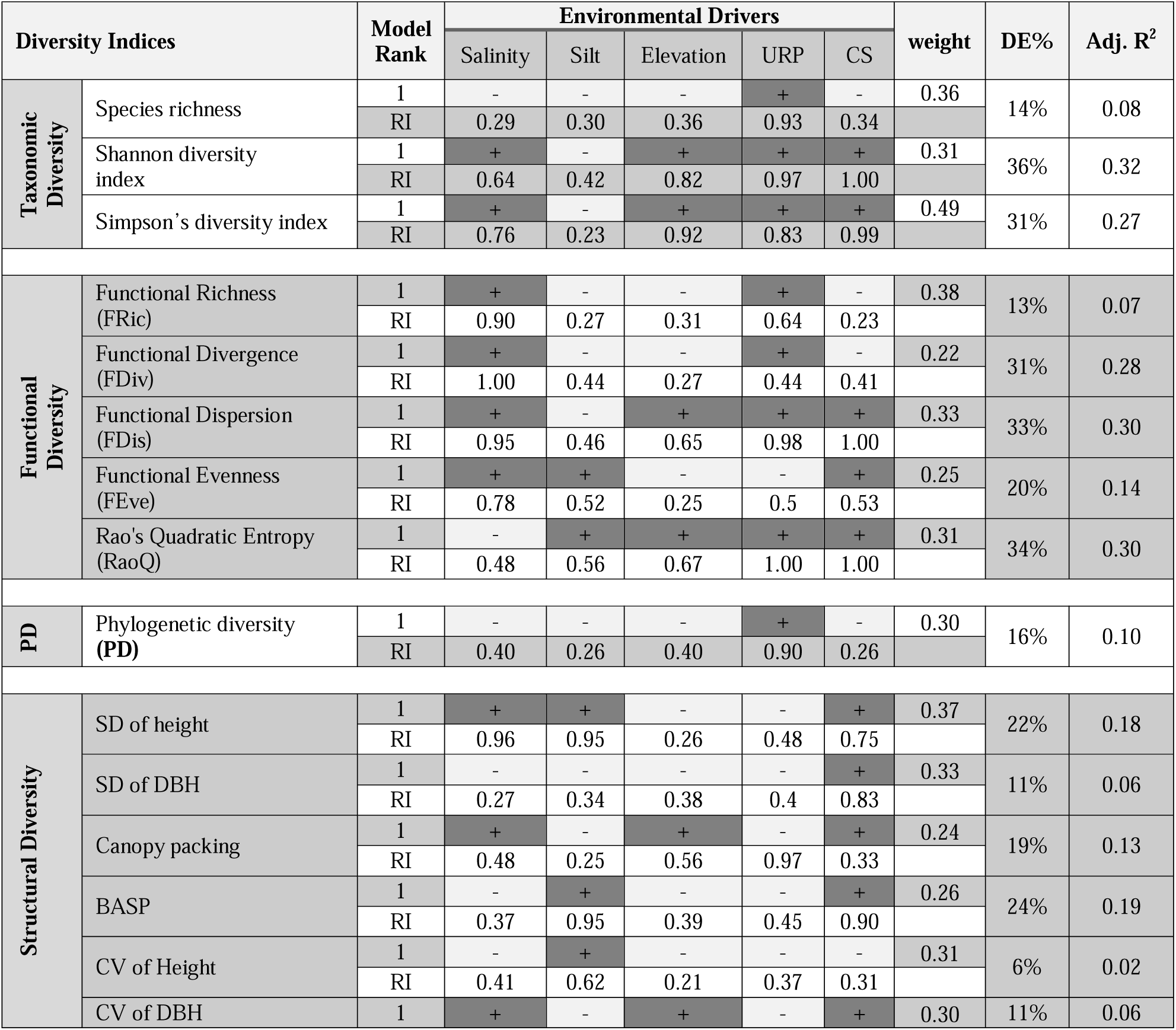
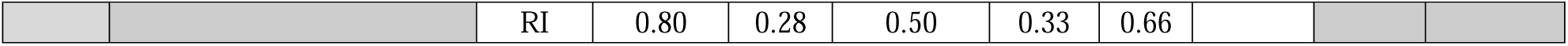
Detail results of Generalized Additive Models (GAMs) for the Taxonomic, Functional and Phylogenetic and Structural Diversity (DE = deviance explained). Only the best model results are included and for the results of all the confidence set models see Table S2.

The relative importance (RI) of the variables in influencing TD, FD, PD and SD diversity indices substantially varied. Salinity, elevation and community structure (CS) had stronger effects on relative-abundance based TD indices (Shannon and Simpson’s diversity) than presence-absence based index (i.e., species richness) while URP (representing the downstream-upstream gradient) showed strong influence on all these TD measures. Salinity, URP and CS were the predominant drivers for the FD indices although siltation had moderate influence on RaoQ (RI=0.56) and FEve (RI=0.52), and elevation showed considerable influence on FDis (RI=0.65) and RaoQ (RI=0.67). While URP showed strongest effects on the spatial distribution of PD, it had least influence on the SD measures. In fact, CS, siltation and salinity were the key drivers for spatial variability in the SD measures.

The partial response plots of the GAMs (Figs. 6 and 7) revealed that all the biodiversity components i.e., TD, FD, PD and SD showed mixed responses to the variables. All the TD indices sharply declined with increasing salinity levels (>7 dS m-1), silt (> 20%) and community structure (>400 tress/0.02 ha) while they all showed an increasing pattern with rising elevation (>2m) and URP (>50 km, upstream areas). In term of FD indices, FRic decreased with increasing URP (>50 km) while increased with increasing salinity (> 8 dS m-1) and elevation (>2.5 m). RaoQ, FDis and FEve showed a decreasing pattern with increasing salinity, URP and CS although their values showed an increasing trend in relatively elevated sites (>2.5 m). PD sharply increased in elevated (>2.5m) upstream areas (URP > 50 km) while decreased with increasing salinity, silt and CS. In terms of SD indices, with increasing salinity SD of height, BASP dropped while canopy packing, SD of DBH, CV of height and CV of DBH increased. Increasing CS and siltation was resulted in decreasing SD of height, SD of DBH, BASP, CV of height and CV of DBH.

**Figure 6.**
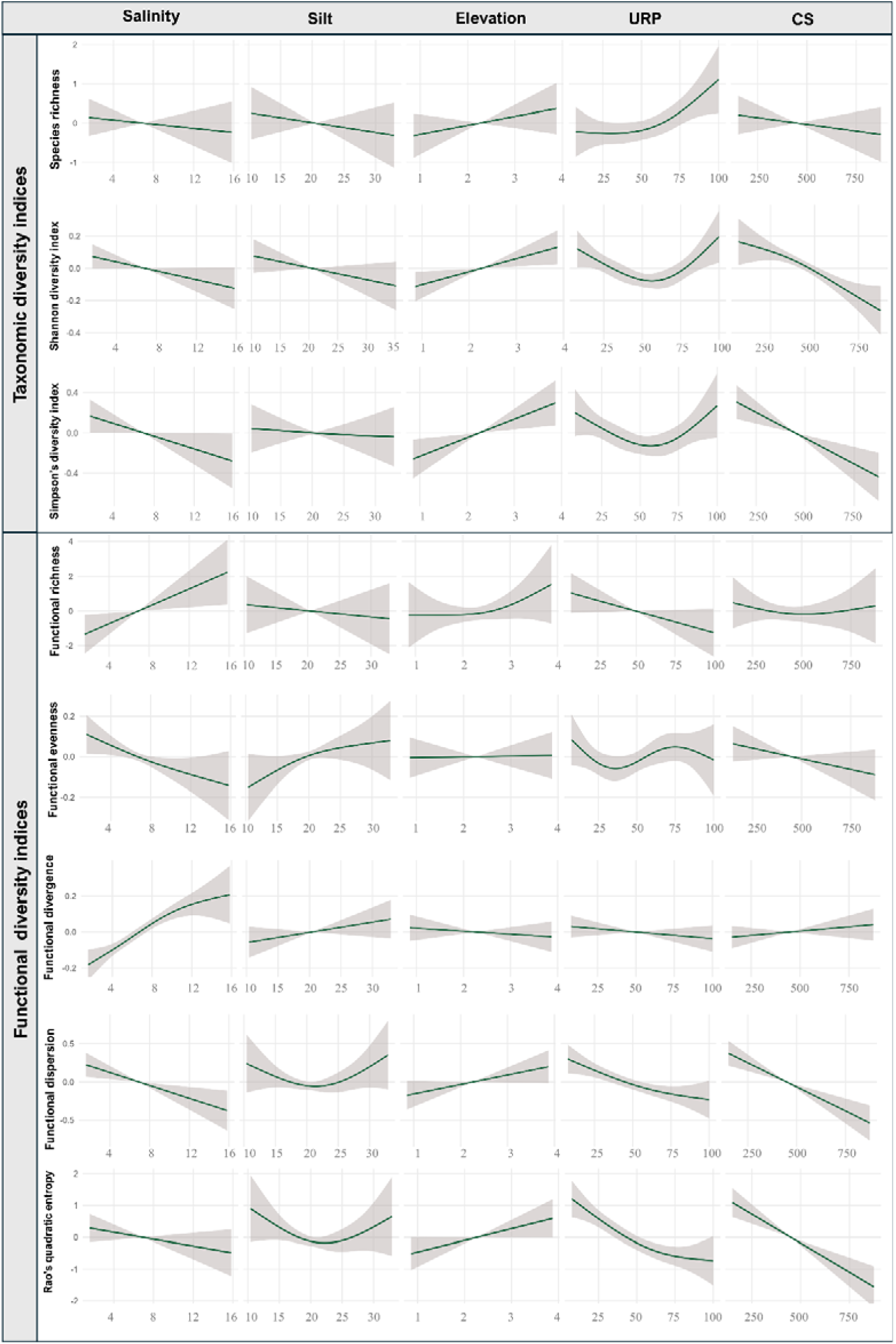
Effects of variables inferred from the best GAMs fitted to the taxonomic and functional diversity indices. The solid line in each plot is the estimated spline function (on the scale of the linear predictor) and shaded areas represent the 95% confidence intervals. Variable units: salinity = dS m^-1^, upriver position = % upriver, elevation = m (above average-sea), community structure = density of all stems for each plot, silt = %.

**Figure 7.**
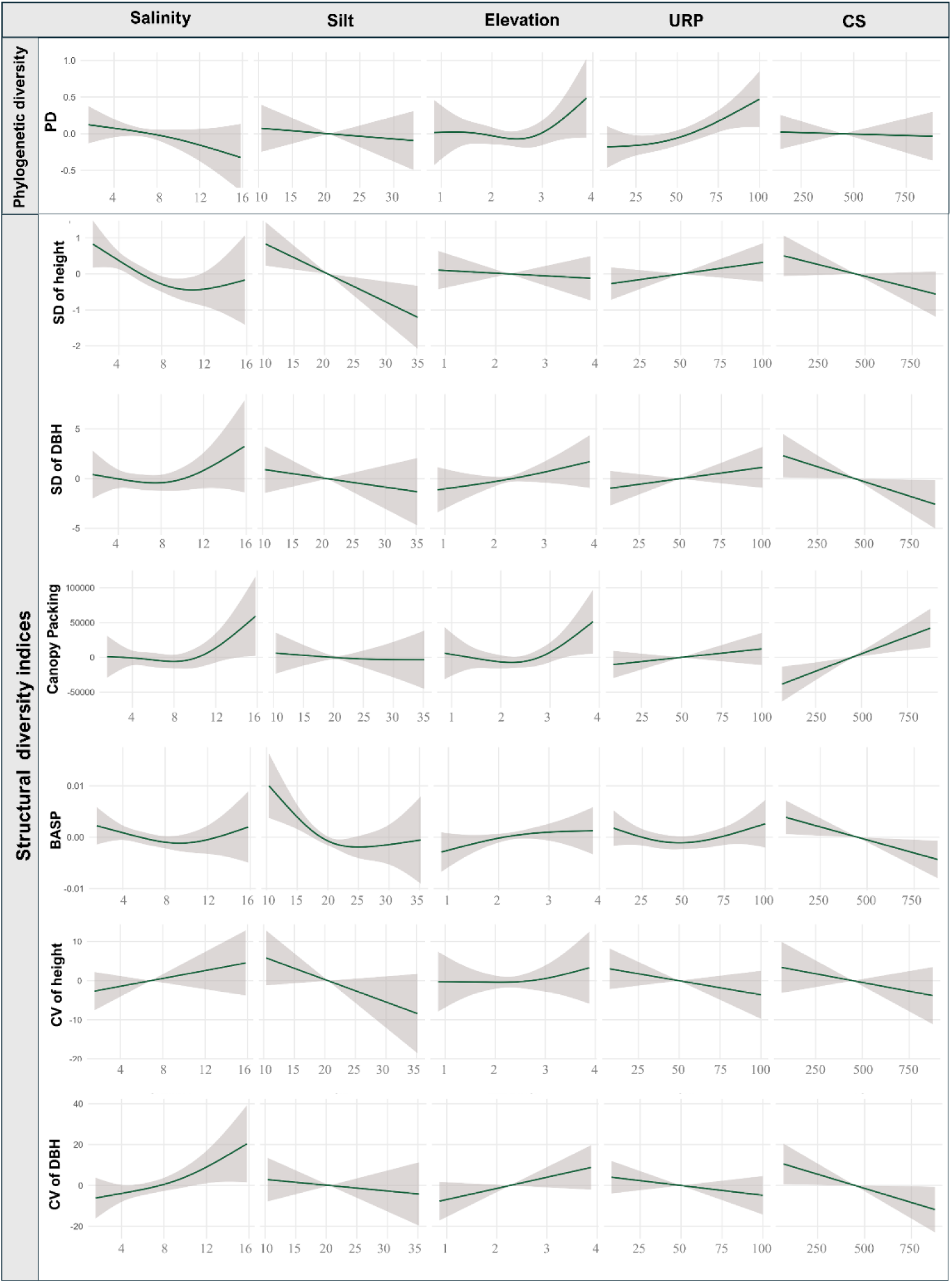
Effects of variables inferred from the best GAMs fitted to the phylogenetic and structural diversity indices. The solid line in each plot is the estimated spline function (on the scale of the linear predictor) and shaded areas represent the 95% confidence intervals. Variable units: salinity = dS m^-1^, upriver position = % upriver, elevation = m (above average-sea), community structure = density of all stems for each plot, silt = %.

### 3.4 Spatial distribution of structural, functional, taxonomic and phylogenetic diversity (Q3)

Spatial structural, taxonomic, functional and phylogenetic diversity maps are presented in Fig. 8. Structural diversity maps uncovered that (i) mangrove communities with greater variations in tree height were mainly distributed in the eastern hyposaline and central mesosaline zones (Fig. 8a); (ii) mangrove communities with greater variations in tree diameter were predominantly confined to the north-western hypersaline zone while diameter distribution is more or less homogeneous across the remaining areas in the Sundarbans (Fig. 8b); (iii) mangrove communities with greater spatial arrangement and density of tree crowns were restricted in some areas (Sarankhola, Harintana, Tambulbunia forest areas) of the eastern hyposaline zone and also in the areas (Koikhali, Kadamtala, Dobeki, Notabeki forest areas) of the north-western hypersaline zone where variations in tree height and DBH were higher than the rest of the ecosystem (Fig. 8c); (iv) northern hyposaline zone comprising the climax mangrove communities had least stand density while stand density hotspots were located in the south-eastern hyposaline zone (Supati, Katka, Kochikhali forest areas) that receives abundant freshwater throughout the year supports (Fig. 8d); (h) similar to stand density, basal area hotspots were located in the south-eastern hyposaline zone (Fig. 8e). Taxonomic diversity maps revealed that species richness (Fig. 8f) and Shannon diversity (Fig. 8g) hotspots were located in the northern hyposaline zone (Kalabogi, Dhangmari, Koira forest areas) while the communities in the western hypersaline zone were least diverse. Phylogenetic diversity map (Fig. 8h) revealed that phylogenetic diversity hotspots were also restricted to the northern hyposaline zone (Kalabogi, Dhangmari, Koira forest areas) with additional distribution in some specific forest areas (Harintana, Tambulbunia forest areas) in the eastern hyposaline zone. Functional diversity maps uncovered that functionally rich mangrove communities were distributed in the north-western hypersaline and sea-dominated southern areas (Fig. 8i) while hypo and mesosaline zones were supporting the mangrove communities with higher functional evenness (Fig. 8k) and taxonomically diverse communities of the hyposaline zone under the Chandpai and Sarankhola Forest rangers had least functional divergence (Fig.8j).

**Figure 8.**
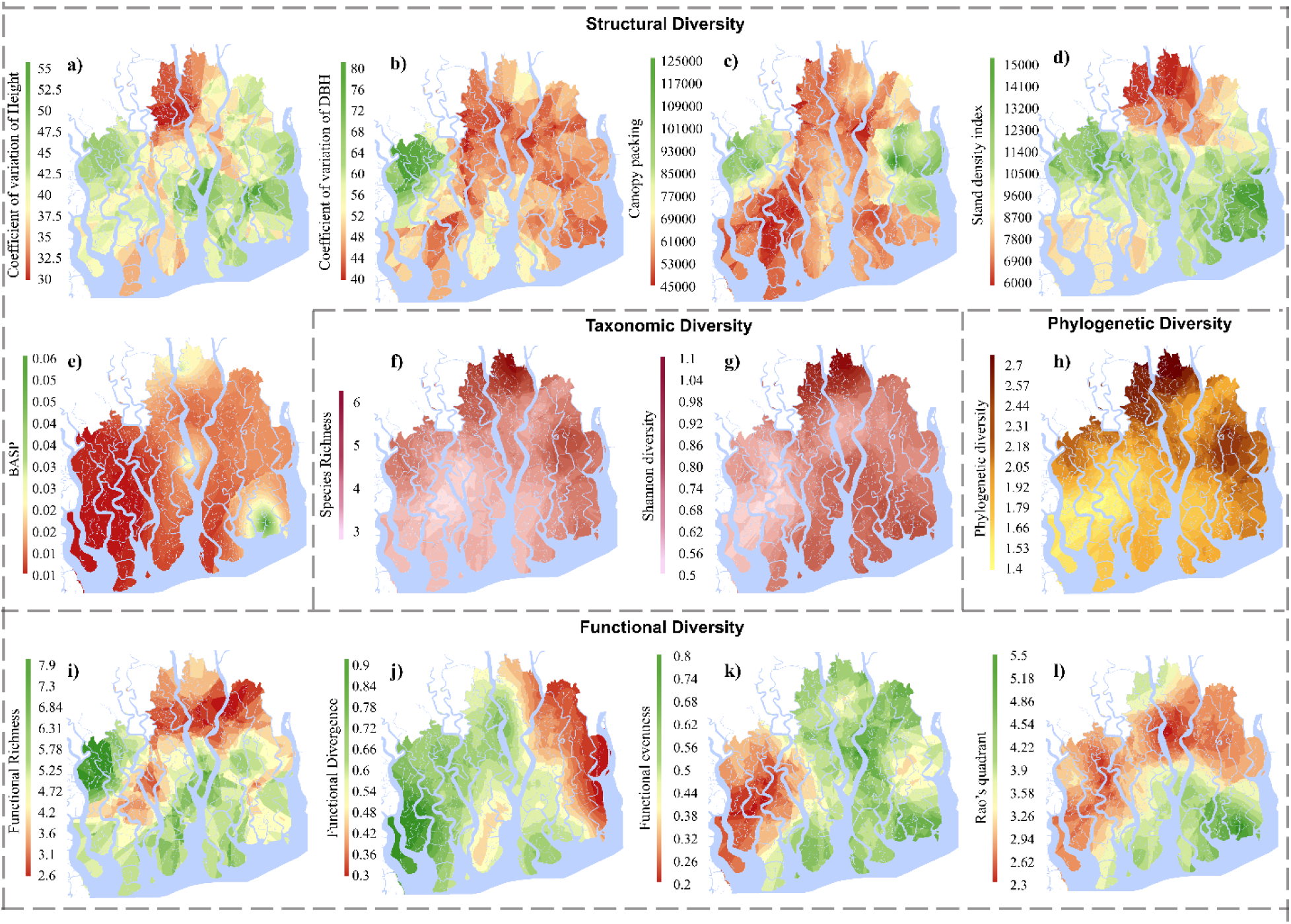
Spatial distribution of mangrove biodiversity dimensions across the Sundarbans, based on data from 110 permanent sample plots. Structural diversity is represented by (a) Coefficient of Variation (CV) of Height, (b) CV of DBH, (c) Canopy Packing, and (d) Stand Density Index. (e) Basal Area per Species (BASP), Taxonomic diversity is depicted using (f) Species Richness, and (g) Shannon Diversity Index. (h) Phylogenetic diversity is represented using Faith’s Phylogenetic Diversity (PD). Functional diversity components include (i) Functional Richness, (j) Functional Divergence, (k) Functional Evenness, and (l) Rao’s Quadratic Entropy.

## 4. DISCUSSION

Our results indicate that the multidimensional aspects of Sundarbans comprising structural, functional, taxonomic, and phylogenetic diversity are interconnected (fig: yet respond differently to environmental stressors (fig: 4) suggesting distinct mechanisms are at play. We also observed that structural diversity metrics showed positive relationships with taxonomic diversity, particularly species richness, and moderate positive relationships with functional diversity (FRic, RaoQ) and phylogenetic diversity. Besides, taxonomic diversity showed positive correlations with functional richness and phylogenetic diversity, indicating close linkages and supporting our first hypothesis. Among environmental variables, salinity showed negative relationships with most structural, taxonomic, and functional diversity metrics, while some functional indices (FRic, FDiv) increased with salinity. Moreover, we observed in GAM results that diversity metrices had mixed and nonlinear responses to salinity, siltation, elevation, upriver proximity, and community structure. In spatial analyses, taxonomic and phylogenetic diversity were higher in northern hyposaline zones (Kalabogi, Dhangmari, and Koira), while functional richness was higher in hypersaline and southern sea-dominated areas, including Koikhali, Kadamtala, Dobeki, Notabeki.

### 4.1 Interconnection of Structural, Functional, Taxonomic, and Phylogenetic Diversities (HI)

We found that structural, functional, taxonomic, and phylogenetic diversity in the Sundarbans are positively interconnected, supporting our first hypothesis. Among structural diversity metrics, standard deviation of height (SD_ht), standard deviation of DBH (SD_DBH), coefficient of variation of height (CV_ht), coefficient of variation of DBH (CV_DBH), and basal area per plot (BASP) showed positive relationships with taxonomic diversity indices, particularly species richness. This finding suggests that mangrove communities with greater variation in tree height and diameter tend to support a larger number of species (Storch et al., 2023). Nevertheless, we found the relationship was weaker with Shannon and Simpson diversity, suggesting that structural diversity is more closely related to species presence than to species relative abundance. Besides, we also found that structural diversity metrics like SD_ht, SD_DBH, CV_ht, CV_DBH, and BASP are correlated positively with functional richness (FRic) and Rao’s quadratic entropy (RaoQ), as well as with phylogenetic diversity (PD), suggesting that structurally heterogeneous communities tend to support broader functional trait diversity and evolutionary lineages (Yuan et al., 2020).

However, our results showed that the stand density index (SDI) maintain moderate negative relationships with most taxonomic diversity indices (species richness, Shannon, Simpson), functional diversity indices (FEve, FDis, RaoQ), and phylogenetic diversity (PD), indicating that higher stand density may be associated with lower multidimensional diversity likely due to increased competition among trees and dominance of a few species (Zhang et al., 2021). In addition to, we observed the strongest correlations between species richness and functional richness (FRic) and between species richness and phylogenetic diversity (PD). This finding suggest that presence of more species in a community allow larger niche space (Song et al., 2014) and a wider range of evolutionary lineages (Zhu et al., 2019). However, their relationship with Shannon and Simpson was weaker, suggesting that functional and phylogenetic diversity are more closely related to species presence rather than relative abundance. Furthermore, functional richness (FRic) and phylogenetic diversity (PD) showed moderate positive relationships, suggesting that communities with greater functional trait diversity also tend to include species from diverse evolutionary backgrounds (Thompson et al., 2015).

### 4.2 Influence of Stressors on Structural, Functional, Taxonomic, and Phylogenetic Diversities (HII)

Our findings show that multiple environmental stressors affect structural, taxonomic, functional, and phylogenetic diversity in the Sundarbans. Among all the environmental variables, salinity had prominent impacts on structural, taxonomic, functional, and phylogenetic diversities displaying consistent negative relationships with several diversity metrics. Particularly with all the taxonomic metrices including species richness, Shannon diversity and Simpson diversity. This finding indicates that with increasing salinity only few salts tolerant species will be able to survive, reducing the evenness of their distribution as well. In the Sundarbans. This pattern occurs because high salinity allows physiological stress for many mangrove species, limiting their growth, site quality and ability to compete for resources (Ahmed et al., 2022b). Similarly functional evenness and dispersion showed negative association with salinity even though functional richness and divergence showed positive. The unexpected increase in functional richness (FRic) and divergence (FDiv) at high salinity (fig: 4) reveals “salinity-driven” trait expansion Due to salinity-driven environmental filtering, this pattern arises where only a few salt-tolerant species *(Excoecaria agallocha, Ceriops decandra)* thrive under high-saline regions (Ahmed et al., 2022b). These species often dominate the community making it homogenous (Sarker, Matthiopoulos, et al., 2019a) and decreasing the even distribution of functional attributes. Furthermore, the remaining species may retain distinct adaptive traits related to salt tolerance, such as specialized leaf structures, salt exclusion mechanisms, or conservative resource-use strategies. As a result, although the number of species decreases and trait distribution becomes less even, the surviving species occupy relatively different functional niches, which increase functional richness and divergence.

Beyond salinity, siltation also had impacts on structural diversity (SD) in the Sundarbans. We found the negative relationships between silt content and the standard deviation of tree height (SD_ht) and basal area per plot (BASP). This finding suggest that high sediment deposition hinder tree growth, favors the expansion of specific species like *Excoecaria* and *Bruguiera* over others (WAHEDUZZAMAN et al., 2022) and lessen variation in canopy structure. As, excessive silt create low-oxygen conditions in the soil around the roots, which makes it difficult for trees to grow properly. Therefore, forests in areas with high silt tend to become more structurally uniform, with trees having similar heights. community structure (CS) also showed similar pattern, which represents tree density. We also observed that increasing CS was associated with lower diversity because dense stands create strong competition (Yang et al., 2024) for light, nutrients, and space.

In contrast, elevation and upriver proximity (URP) has lower effect on diversities than the others. We observed elevation (above ∼2.5 m) and URP is positively correlated with phylogenetic diversity (PD) suggesting that slightly higher upstream areas may support species from more diverse evolutionary lineages. As these areas are less frequently flooded by tides and experience lower salt accumulation, creating more favorable conditions for different species to coexist. However, URP had lower effect on structural diversity (SD) (fig: 5), suggesting that while the evolutionary composition of species may change along the river gradient, the physical structure of the forest is mainly shaped by salinity, siltation and tree density.

### 4.3 Spatial Patterns in Structural, Functional, Taxonomic, and Phylogenetic Diversities (HIII)

Our results show distinct spatial patterns across the Sundarbans in different saline zones. We found that species richness, Shannon diversity, and phylogenetic diversity (PD) were higher in the northern hyposaline areas, such as Dhangmari and Kalabogi. As these areas receive freshwater and experience lower salinity which allows for larger number of species and diverse evolutionary lineages to coexist. However, in western hypersaline zone, they were lower. Due to high salinity which acts as a barrier that restricts species survival. However, we observed noticeable patterns for functional and structural diversity. The north-western hypersaline zone and southern coastal areas showed high functional richness (FRic). This finding indicate that even though fewer species occur in these areas, the species that remain have different functional traits that help them survive under stressful conditions such as high salinity and strong tidal influence.

For structural diversity, the south-eastern hyposaline areas, such as Katka and Kochikhali, showed high stand density and basal area (BASP). Due to better freshwater input and higher productivity in these areas. In addition, our results show that higher variation in tree height (CV of height) appeared mainly in the eastern and central zones. This result is also due to availability of freshwater in those areas. These distinct spatial patterns reveal that the Sundarbans is a mosaic of specialized ecological zones, where taxonomic richness does not always overlap with functional or structural uniqueness, emphasizing the need for multidimensional strategies for effective conservation.

### 4.4 Research Implications and Future Directions

The findings of this study shed critical insights into how the multidimensionality of biodiversity regulates the resilience of the Sundarbans under changing conditions. We observe that in hyposaline regions, the strong connection between taxonomic and structural diversity forms a “stability anchor,” where high species richness and vertical complexity coexist to maximize resource use and biomass accumulation. This suggests that maintaining freshwater flow is not about preserving species counts only, but about protecting the structural integrity that allows these forests to function as high-capacity carbon sinks. Conversely, in hypersaline and sea-dominated zones, we identify a “functional compensation” mechanism. Despite a decline in taxonomic richness, the increase in functional richness (FRic) and divergence (FDiv) suggests that specialized halophytes occupy unique niches to maintain ecosystem processes under extreme stress. This indicates that in the face of sea-level rise, the survival of the forest may depend on these functionally distinct species. Protecting these “stress-tolerant hotspots” in the western and southern Sundarbans is therefore essential, as they possess those specific traits which are required to maintain a protective coastal shield. Furthermore, the “structural stagnation” and reduced phylogenetic diversity observed in high-silt and high-salinity areas reflects a significant vulnerability to climate change. As these stressors intensify, the resulting homogenization of the canopy (reduced CV of height) may diminish the forest’s ability to attenuate tidal energy during cyclonic events. Given the frequent mortality in pioneer stands and the formation of gaps following natural calamities, our research suggests that a multidimensional management approach is required. Restoration efforts should focus on “trait-based enrichment,” where a mix of species with diverse evolutionary lineages and structural forms are introduced into gaps to expedite recovery and enhance long-term resilience. This research holds strong applicability for global mangrove conservation and nature-based solutions. By moving beyond single-dimension monitoring, management can better prioritize areas that provide the highest co-benefits of carbon sequestration, structural protection, and evolutionary preservation. It underscores the necessity of a holistic framework to fortify coastal belts against the synergistic challenges of hyper salinity, sedimentation, and climate-driven disturbances.

Although the results of this study provide a fundamental understanding of multidimensional approach in mangroves, primary location of this study can be generalized, and the findings might not be as applicable to other mangrove systems with distinct geophysical environments. To validate these patterns on a larger scale, comparative research across several geographical sites would be necessary. Additionally, the temporal interplay between seasonal rainfall patterns, water depth, hydrological parameters and varying salinity levels was not taken into consideration in our dataset. Over time, species’ physiological reactions and community formation are probably influenced by these dynamic hydrological conditions. Future research incorporating a broader range of functional traits and socio-ecological factors would provide a more holistic systems perspective. Such an approach is crucial to translate these ecological findings into feasible, high-impact conservation and restoration strategies.

## 5. CONCLUSIONS

This study analyzed the interconnection among structural, functional, taxonomic and phylogenetic diversities and how they respond to multiple stressors in the Sundarbans. Our results revealed that mangrove diversities are substantially affected by salinity and siltation. We also found strong relationship between structural and taxonomic diversity. However, functional and phylogenetic diversity showed distinct patterns, with some metrics rising in high-salinity areas where stress-tolerant species maintain unique ecological roles. As species richness is often used as a proxy for other types of diversity, our findings prove that structural, functional and diverse evolutional lineages provide a more holistic approach for forest health. These findings are crucial for advancing global mangrove conservation, expanding our understanding of ecosystem resilience, and informing management strategies that protect both ecological and evolutionary history in coastal forests like Sundarbans.

## CRediT authorship contribution statement

Conceptualization-SKS (lead), BD. Data Collection and organization- SKS (lead), NKP, AAS, HAMF, ZBM, MAU, ZBM, AMZ, MMHK. Data analysis- BD (lead), SKS, AAS. Manuscript writing- BD (lead), SKS, SA, HX, EJE. Review and Editing- SKS (lead), SA, HX, NKP, IA, EJE, AAS, MMHK, MAU. Final review- SKS, SA, HX, HAMF, ZBM, and IA. All authors read and approved the final version of the manuscript.

## Acknowledgements

We sincerely acknowledge and thank the Bangladesh Forest Department for providing all logistic support during the fieldwork. SKS acknowledges the supports of the University of Glasgow, SUST Research Centre (Project IDs: FES/2024/1/03, FES/2023/2/08, FES/2025/1/03) and the Commonwealth Scholarship Commission, United Kingdom. An earlier version of this manuscript formed a chapter of BD’s MSc thesis.

## Appendix A. Supplementary data

Supplementary Materials (.docx)

## Data availability

Data and scripts will be made available in Bangladesh Forest Department’s Bangladesh Forest Information System (http://bfis.bforest.gov.bd/bfis/) and can be accessed upon permission.

