## Supplemental Table 1 for "Biodiversity Dimensions in Mangroves: Uncovering Interactions and Spatial Drivers in the Sundarbans"

**Title**

**Running head**

Biodiversity Dimensions in Mangroves

**Authors**

Bornali Das^1^, Abdullah Al Asif^1^, Shamim Ahmed^2^, Huang Xingyun^3^, H. A. M. Fayeem^4^, Zawyad Bin Mostofa^4^, Esrat Jahan Ema^5^, Adel Mahmud Zaddary^6^, Md Amanat Ullah^7^, Md. Mehedi Hasan Khan^8^, Nirmal Kumar Paul^9^, Imran Ahmed^10^, Swapan Kumar Sarker^1*^

**Affiliation**

^1^ Department of Forestry & Environmental Science, Shahjalal University of Science & Technology, Sylhet 3114, Bangladesh.

^2^ Department of Forest Bioeconomy and Technology, Swedish University of Agricultural Sciences, Vallvägen 9 C, 75007 Uppsala, Sweden.

^3^ South China Botanical Garden, Chinese Academy of Sciences, Xingke Road 723, Guangzhou, China. 519650

^4^ Arannayk Foundation (Bangladesh Tropical Forest Conservation Foundation), 572/K, Wasi Tower, ECB Chattar, Matikata, 1206 Dhaka, Bangladesh.

^5^ Bangladesh Country Office, IUCN (International Union for Conservation of Nature), Banani, Dhaka 1213, Bangladesh.

^6^ Center for Natural Resource Studies (CNRS), Banani, Dhaka 1213, Bangladesh.

^7^ Ecology, Forestry and Biodiversity Division, Center for Environmental and Geographic Information Services (CEGIS), Dhaka-1207, Bangladesh.

^8^ RIMS Unit, Bangladesh Forest Department, Ban Bhaban, Agargaon, Dhaka, Bangladesh.

^9^ Wildlife Management and Nature Conservation Division, Khulna, Bangladesh Forest Department, Bangladesh.

^10^ Khulna Circle, Bangladesh Forest Department, Khulna.

Supplementary Figures

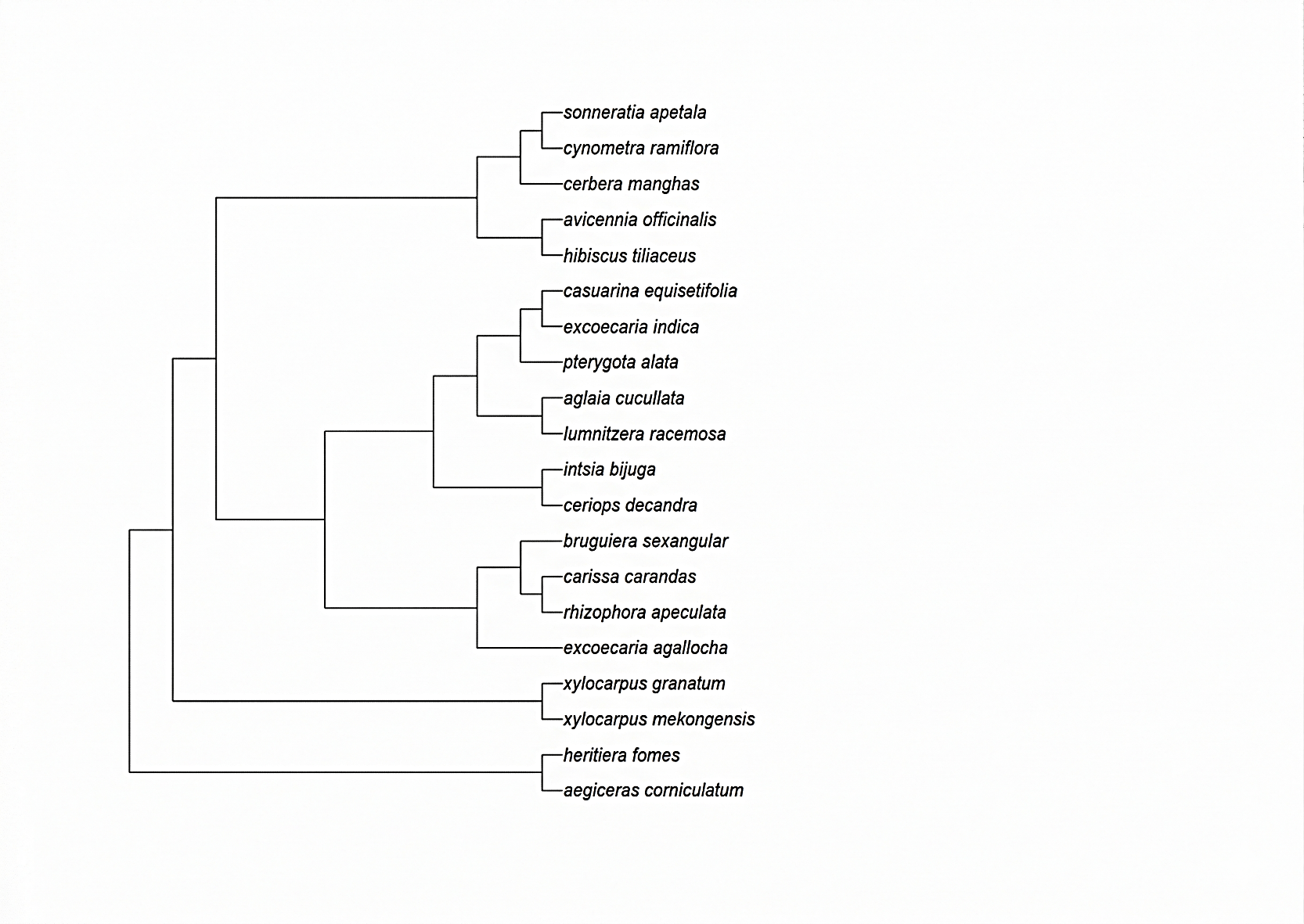

**Figure S1.** Phylogenetic tree of mangrove plant species recorded in the Sundarbans mangrove forest used for the calculation of phylogenetic diversity (PD). The dendrogram illustrates the evolutionary relationships among the species included in the study. This phylogenetic framework was used to quantify plot-level phylogenetic diversity across the 110 permanent sample plots.

Supplementary Tables

Table S1. Descriptive statistics of structural, taxonomic, functional, and phylogenetic diversity metrics across the 110 permanent sample plots in the Sundarbans mangrove forest. The table presents the minimum (Min), maximum (Max), mean, median, range, standard deviation (SD), and coefficient of variation (CV) for each diversity metric.

| Variable | | Min | Max | Mean | Median | Range | SD | CV |
| --- | --- | --- | --- | --- | --- | --- | --- | --- |
| Structural diversity metrics | Height | 1.10 | 25.00 | 6.29 | 5.50 | 122.2 | 10.26 | 0.81 |
|  | DBH | 1.80 | 124.00 | 12.71 | 10.00 | 23.9 | 3.79 | 0.60 |
|  | Stand Density Index | 1700.00 | 21875.00 | 11229.32 | 10987.50 | 20175.00 | 4493.67 | 0.40 |
|  | Canopy Packing | 5231.46 | 381729.22 | 77830.72 | 61526.90 | 376497.75 | 59525.59 | 0.76 |
|  | Basal Area Per Plot | 0.01 | 0.08 | 0.02 | 0.02 | 0.07 | 0.01 | 0.46 |
| Taxonomic diversity metrics | Species Richness | 2 | 1 | 4 | 4 | 8 | 1.48 | 0.35 |
|  | Shannon diversity index | 0.05 | 1.51 | 0.75 | 0.73 | 1.46 | 0.27 | 0.36 |
|  | Simpson diversity index | 1.02 | 4.08 | 1.88 | 1.81 | 3.06 | 0.53 | 0.28 |
| Functional diversity  metrics | Functional Richness | 0.47 | 15.68 | 5.27 | 4.83 | 15.21 | 3.40 | 0.64 |
|  | Functional Evenness | 0.05 | 0.98 | 0.53 | 0.50 | 0.93 | 0.25 | 0.48 |
|  | Functional Divergence | 0.23 | 1.00 | 0.71 | 0.73 | 0.77 | 0.20 | 0.28 |
|  | Functional Dispersion | 0.07 | 3.30 | 1.66 | 1.78 | 3.23 | 0.54 | 0.33 |
|  | Rao’s Quadratic Entropy | 0.15 | 11.84 | 3.86 | 3.89 | 11.69 | 1.48 | 0.38 |
| Phylogenetic diversity (Faith’s PD) | | 1.53 | 5.16 | 3.30 | 3.42 | 3.63 | 0.75 | 0.23 |

Table S2 Detailed results of the Generalized Additive Models (GAMs) explaining patterns of structural, functional, taxonomic, and phylogenetic diversity across the study plots. The table includes the deviance explained (DE) by each model and the relative importance (RI) of predictor variables. The environmental covariates include salinity, silt, elevation, upriver position (URP), and community size (CS). Here, RI (Relative Importance) represents the contribution of each predictor variable to the fitted GAMs based on model selection procedures.

| Structural Diversity | | | | | | | | | | | |
| --- | --- | --- | --- | --- | --- | --- | --- | --- | --- | --- | --- |
| Diversity  Indicators | Model Rank | Variables | | | | | | | |  |  |
|  |  | Salinity | Silt | Elevation | URP | CS | df | delta | weight | DE% | R-sq.(adj) |
| SD  Of  height | 1 | + | + | - | - | + | 6 | 0 | 0.37 | 22% | 0.18 |
|  | 2 | + | + | - | + | + | 7 | 0.73 | 0.26 |  |  |
|  | 3 | + | + | - | + | - | 6 | 1.90 | 0.14 |  |  |
|  | 4 | + | + | + | - | + | 7 | 2.15 | 0.12 |  |  |
|  | 5 | + | + | + | + | + | 8 | 2.90 | 0.08 |  |  |
|  | RI | 0.96 | 0.95 | 0.26 | 0.48 | 0.75 |  |  |  |  |  |
| SD  Of  DBH | 1 | - | - | - | - | + | 3 | 0 | 0.33 | 11% | 0.06 |
|  | 2 | - | - | + | - | + | 4 | 1.03 | 0.20 |  |  |
|  | 3 | - | - | - | + | + | 4 | 1.21 | 0.18 |  |  |
|  | 4 | - | + | - | - | + | 4 | 1.51 | 0.16 |  |  |
|  | 5 | + | - | - | - | + | 5 | 1.96 | 0.13 |  |  |
|  | RI | 0.27 | 0.34 | 0.38 | 0.4 | 0.83 |  |  |  |  |  |
| Canopy  packing | 1 | + | - | + | - | + | 7 | 0 | 0.24 | 19.2% | 0.13 |
|  | 2 | - | - | + | - | + | 6 | 0.08 | 0.24 |  |  |
|  | 3 | - | - | - | - | + | 3 | 0.25 | 0.22 |  |  |
|  | 4 | + | - | - | - | + | 5 | 0.86 | 0.16 |  |  |
|  | 5 | + | - | + | + | + | 9 | 1.06 | 0.14 |  |  |
|  | RI | 0.48 | 0.25 | 0.56 | 0.97 | 0.33 |  |  |  |  |  |
| BASP | 1 | - | + | - | - | + | 5 | 0 | 0.26 | 24% | 0.19 |
|  | 2 | - | + | - | + | + | 8 | 0.20 | 0.23 |  |  |
|  | 3 | - | + | + | - | + | 7 | 0.63 | 0.19 |  |  |
|  | 4 | + | + | - | - | + | 7 | 0.76 | 0.18 |  |  |
|  | 5 | - | + | + | + | + | 9 | 1.15 | 0.15 |  |  |
|  | RI | 0.37 | 0.95 | 0.39 | 0.45 | 0.9 |  |  |  | 6% | 0.02 |
| CV of Height | 1 | - | + | - | - | - | 3 | 0 | 0.31 |  |  |
|  | 2 | + | + | - | - | - | 4 | 0.80 | 0.21 |  |  |
|  | 3 | - | + | - | - | + | 4 | 1.02 | 0.18 |  |  |
|  | 4 | - | - | - | - | - | 2 | 1.13 | 0.17 |  |  |
|  | 5 | + | - | - | - | - | 3 | 1.70 | 0.13 |  |  |
|  | RI | 0.41 | 0.62 | 0.21 | 0.37 | 0.31 |  |  |  |  |  |
| CV of DBH | 1 | + | - | + | - | + | 6 | 0 | 0.30 | 11% | 0.06 |
|  | 2 | + | - | - | - | + | 5 | 0.28 | 0.26 |  |  |
|  | 3 | + | - | + | + | + | 6 | 1.22 | 0.16 |  |  |
|  | 4 | + | - | - | - | - | 3 | 1.36 | 0.15 |  |  |
|  | 5 | + | - | - | + | + | 5 | 1.56 | 0.14 |  |  |
|  | RI | 0.8 | 0.28 | 0.5 | 0.33 | 0.66 |  |  |  |  |  |
| Functional Diversity | | | | | | | | | | | |
| Diversity  Indicators | Model Rank | Variables | | | | | | | |  |  |
|  |  | Salinity | Silt | Elevation | URP | CS | df | delta | weight | DE% | R-sq.(adj) |
| Functional Richness | 1 | + | - | - | + | - | 4 | 0 | 0.38 | 13% | 0.07 |
|  | 2 | + | - | - | - | - | 3 | 1.21 | 0.21 |  |  |
|  | 3 | + | - | + | + | - | 6 | 1.68 | 0.16 |  |  |
|  | 4 | + | + | - | + | - | 5 | 1.97 | 0.14 |  |  |
|  | 5 | + | - | - | + | + | 5 | 2.48 | 0.11 |  |  |
|  | RI | 0.9 | 0.27 | 0.31 | 0.64 | 0.23 |  |  |  |  |  |
| Functional Divergence | 1 | + | - | - | + | - | 5 | 0 | 0.22 | 31% | 0.28 |
|  | 2 | + | - | - | - | - | 4 | 0.00 | 0.22 |  |  |
|  | 3 | + | - | - | - | + | 5 | 0.07 | 0.21 |  |  |
|  | 4 | + | + | - | - | - | 5 | 0.28 | 0.19 |  |  |
|  | 5 | + | + | - | + | - | 6 | 0.43 | 0.17 |  |  |
|  | RI | 1 | 0.44 | 0.27 | 0.44 | 0.41 |  |  |  |  |  |
| Functional Dispersion | 1 | + | - | + | + | + | 6 | 0 | 0.33 | 33% | 0.3 |
|  | 2 | + | + | + | + | + | 9 | 0.30 | 0.29 |  |  |
|  | 3 | + | - | - | + | + | 5 | 0.92 | 0.21 |  |  |
|  | 4 | + | + | - | + | + | 8 | 1.60 | 0.15 |  |  |
|  | 5 | - | + | + | + | + | 8 | 5.31 | 0.02 |  |  |
|  | RI | 0.95 | 0.46 | 0.65 | 0.98 | 1 |  |  |  |  |  |
| Functional Evenness | 1 | + | + | - | - | + | 5 | 0 | 0.25 | 20% | 0.14 |
|  | 2 | + | - | - | - | + | 4 | 0.36 | 0.21 |  |  |
|  | 3 | + | - | - | - | - | 3 | 0.37 | 0.21 |  |  |
|  | 4 | + | + | - | - | - | 4 | 0.46 | 0.20 |  |  |
|  | 5 | + | - | - | + | - | 5 | 1.18 | 0.14 |  |  |
|  | RI | 0.78 | 0.52 | 0.25 | 0.5 | 0.53 |  |  |  |  |  |
| Rao's Quadratic Entropy | 1 | - | + | + | + | + | 8 | 0 | 0.31 | 34% | 0.3 |
|  | 2 | + | - | + | + | + | 7 | 0.76 | 0.21 |  |  |
|  | 3 | + | + | + | + | + | 9 | 0.97 | 0.19 |  |  |
|  | 4 | - | - | + | + | + | 6 | 1.53 | 0.15 |  |  |
|  | 5 | - | + | - | + | + | 7 | 1.76 | 0.13 |  |  |
|  | RI | 0.48 | 0.56 | 0.67 | 1 | 1 |  |  |  |  |  |
| Taxonomic Diversity | | | | | | | | | | | |
| Diversity  Indicators | Model Rank | Variables | | | | | | |  |  |  |
|  |  | Salinity | Silt | Elevation | URP | CS | df | delta | weight | DE% | R-sq.(adj) |
| Species richness | 1 | - | - | - | + | - | 4 | 0 | 0.36 | 14% | 0.08 |
|  | 2 | - | - | + | + | - | 5 | 1.29 | 0.19 |  |  |
|  | 3 | - | - | - | + | + | 5 | 1.59 | 0.16 |  |  |
|  | 4 | - | + | - | + | - | 5 | 1.67 | 0.15 |  |  |
|  | 5 | + | - | - | + | - | 5 | 1.82 | 0.14 |  |  |
|  | RI | 0.29 | 0.3 | 0.36 | 0.93 | 0.34 |  |  |  |  |  |
| Shannon diversity index | 1 | + | - | + | + | + | 8 | 0 | 0.31 | 36% | 0.32 |
|  | 2 | + | + | + | + | + | 9 | 0.30 | 0.27 |  |  |
|  | 3 | - | - | + | + | + | 6 | 0.82 | 0.21 |  |  |
|  | 4 | - | + | + | + | + | 8 | 1.74 | 0.13 |  |  |
|  | 5 | + | - | - | + | + | 6 | 2.61 | 0.08 |  |  |
|  | RI | 0.64 | 0.42 | 0.82 | 0.97 | 1 |  |  |  |  |  |
| Simpson’s diversity index | 1 | + | - | + | + | + | 7 | 0 | 0.49 | 31% | 0.27 |
|  | 2 | - | - | + | + | + | 6 | 1.88 | 0.19 |  |  |
|  | 3 | + | + | + | + | + | 8 | 2.36 | 0.15 |  |  |
|  | 4 | + | - | + | - | + | 5 | 2.69 | 0.13 |  |  |
|  | 5 | - | + | + | + | + | 8 | 4.70 | 0.05 |  |  |
|  | RI | 0.76 | 0.23 | 0.92 | 0.83 | 0.99 |  |  |  |  |  |
| Phylogenetic Diversity | | | | | | | | | | | |
| Diversity  Indicators | Model Rank | Variables | | | | | | | |  |  |
|  |  | Salinity | Silt | Elevation | URP | CS | df | delta | weight | DE% | R-sq.(adj) |
| Phylogenetic  diversity | 1 | - | - | - | + | - | 4 | 0 | 0.30 | 16% | 0.1 |
|  | 2 | - | - | + | + | - | 6 | 0.45 | 0.24 |  |  |
|  | 3 | + | - | - | + | - | 5 | 0.65 | 0.22 |  |  |
|  | 4 | + | - | + | + | - | 7 | 1.61 | 0.13 |  |  |
|  | 5 | - | + | - | + | - | 5 | 1.95 | 0.11 |  |  |
|  | RI | 0.4 | 0.26 | 0.4 | 0.9 | 0.26 |  |  |  |  |  |
